# Electric field intensity modulates keratocyte migration without altering turning dynamics

**DOI:** 10.1101/2025.10.29.684651

**Authors:** Niloofar Pishkari, Eloina Corradi, Gawoon Shim, Daniel Cohen, Marine Luciano, Sylvain Gabriele, Giovanni Cappello, Thomas Boudou, Martial Balland

## Abstract

Cell migration is a cornerstone of biological systems, enabling organisms to adapt to environmental stimuli and maintain homeostasis. Disruptions in this process can lead to functional impairment or system failure. In many cases, cells do not move randomly; instead, they migrate directionally in response to external cues, allowing them to perform essential biological functions. This directed movement is especially important in processes such as morphogenesis, cancer invasion, and wound healing. To unravel the complexities of directional cell migration, investigating natural guiding stimuli is crucial. Among these, electrical fields stand out as precise and physiologically relevant stimulus. Using a platform designed to apply programmable electric fields, the SCHEEPDOG device; we applied controlled electric field of varying intensities to keratocytes and quantitatively analyzed their migratory behavior. Our findings reveal that electric field stimulation not only induces robust directional migration but also enhances migration speed in an intensity-dependent manner. Additionally, cells initially moving in random directions gradually align with the field vector, with higher intensities accelerating the alignment. Intriguingly, while both speed and alignment time can be modulated through stimulation, the overall shape of migration trajectories remains unchanged. In other terms, for cells initially moving to the opposite direction of the field, the alignment is accompanied by making a turn and the size and shape of this turn is not affected by the magnitude of the electrical stimulation. Together, these results demonstrate that electrical stimulation can tune the speed and directional alignment of keratocyte migration without altering turning dynamics. These findings contribute to a deeper understanding of electrotaxis and offers new insights into how biophysical cues regulate cell migration in both physiological and pathological contexts.

## Introduction

Directed cell migration is a fundamental process in multicellular organisms, essential in various physiological and pathological contexts, including embryogenesis^1–3^, wound, immune responses^6,7^, and cancer invasion^8,9^. Given its critical role, the ability to guide or control cell movement presents a powerful tool for both research and therapeutic applications.

Cell migration can be directed through various mechanisms inherent in biological systems, such as chemotaxis^10,11^, durotaxis^12^, haptotaxis^11,13^ and electrotaxis or galvanotaxis (first described by Du Bois-Reymond in 1849)^14^. The latter, allows for the precise manipulation and direction of cell migration in response to electric fields^15,16^.

The process of galvanotaxis is of significant biological importance, and to unravel the mechanisms underlying this phenomenon, a wide range of experimental and theoretical studies have been conducted. Collectively, these investigations have shown that galvanotaxis is a complex, multi-layered process driven by the interplay of various mechanisms. Experimental studies have demonstrated the involvement of several signaling pathways^17–19^, the polarization of charged elements within the cell membrane^20,21^, and changes in membrane potential^22,23^. Theoretical work has contributed by modeling processes such as osmotic pressure modulation across the membrane^24^, offering insight into the physical forces involved in cell migration. Other research has revealed roles for ion channel regulation^25^, reorganizing proteins at cell surface level^26^, and cytoskeletal dynamics ^27^, further emphasizing the integrative nature of this phenomenon. Collectively all these factors govern directional movement and responses to external cues^21,28^. Probably due to the different mechanisms acting in concert, endogenous electric fields in biological systems vary widely in strength, with different studies reporting different ranges of intensities^4,29–31^.

As reported by^32^, many cell types^33–35^ exhibit directed migration across a wide range of electric field strengths and electrical stimulation increases the migration speed of cells^16,21^.

Here, we hypothesize that cell migratory behavior is influenced by electric field intensity. Therefore, we investigate how modulating electric field strength can regulate cell trajectories and guide their migrating trajectories. To address this, we utilized the specialized “SCHEEPDOG” device^16^, which allowed us to reduce the influence of environmental factors such as heat and pH by maintaining a constant flow of media. However, reference to previous works^21^, we know that generated heat by electric fields at high intensities partially can affect the speed of cells, but there is no evidence that directionality is also affected by this factor. Cells exhibited directionality at all electric field intensities, consistent with previous studies^36^. With our approach to tune the electric field intensity, we observed that by increasing electric field intensity, keratocyte cells migrated more rapidly in the direction of the field, covering a greater distance within the same period. Moreover, at higher electric field intensities, the cells show a rapid increase in speed along the field’s direction, reaching their maximum velocity within a few minutes. We also examined the alignment behavior of the cells under varying stimulation conditions. Our findings indicate that as stimulation intensity increases, the time required for cells to align with the electric field decreases, while their speed increases. Interestingly, the travelled distance required for alignment remains constant across different field intensities. When categorizing cells based on their initial migration direction, we observed that cells moving at smaller angles to the electric field align more quickly. Conversely, cells migrating opposite to the field direction take longer to align and cover a greater distance. We further examined the turning behavior of the cells during alignment. While the angular velocity increases with stronger stimulation, the curvature of their migration paths remains largely unaffected by electric field intensity.

Overall, our data demonstrate that both the speed and direction of cell migration can be controlled and fine-tuned by adjusting the intensity of electric field stimulation but the shape of the turn is not dependent on the field strength and might be an intrinsic property of the cells themselves.

## Results

### Electric field intensity regulates keratocyte migration direction and speed

To explore and manipulate cellular migration in a controlled environment, mechanism such as electrotaxis is useful. To study these processes in a controlled environment, we utilize the Spatiotemporal Cellular Herding with Electrochemical Potentials to Dynamically Orient Galvanotaxis (SCHEEPDOG) system^16^ (Fig.1A). This device is connected to a pumping system that continuously provides fresh media for the cells while also removing consumed media and byproducts generated by the Ag/AgCl electrodes, which create the electric field. By linking the system to a source meter, the magnitude of the electric field is controlled precisely, enabling us to stimulate the cells in a reproducible manner (Fig.1A). For our experiments, we used fish epithelial keratocytes isolated from the scales from the Central American cichlid Hypsophrys Nicaraguensis. Keratocytes were seeded onto a clean glass coverslip (Fig.1B), which was then mounted on the SCHEEPDOG platform to be exposed to electric fields for stimulation. Under control conditions, in the absence of electrical stimulation, keratocytes migrate randomly in all directions (Fig.1C, control), displaying a range of migration speeds influenced by environmental factors such as pH^21^ and temperature^37^, closely associated with the adopted cell shape^38^. The random migration without electric field highlight that the flow of medium in the Galvano-tactic system does not influence per se cell direction (Fig.1C, control). However, upon the application of a 2mA electrical field, the cells exhibit a clear directional movement towards the field, as indicated by the negative values along the X-axis in our setup (Fig.1C, EF: 2mA). To further explore the effects of electrical stimulation on cellular behavior, we increased the field intensities to 4, 6, and 8mA.

**Figure 1.**
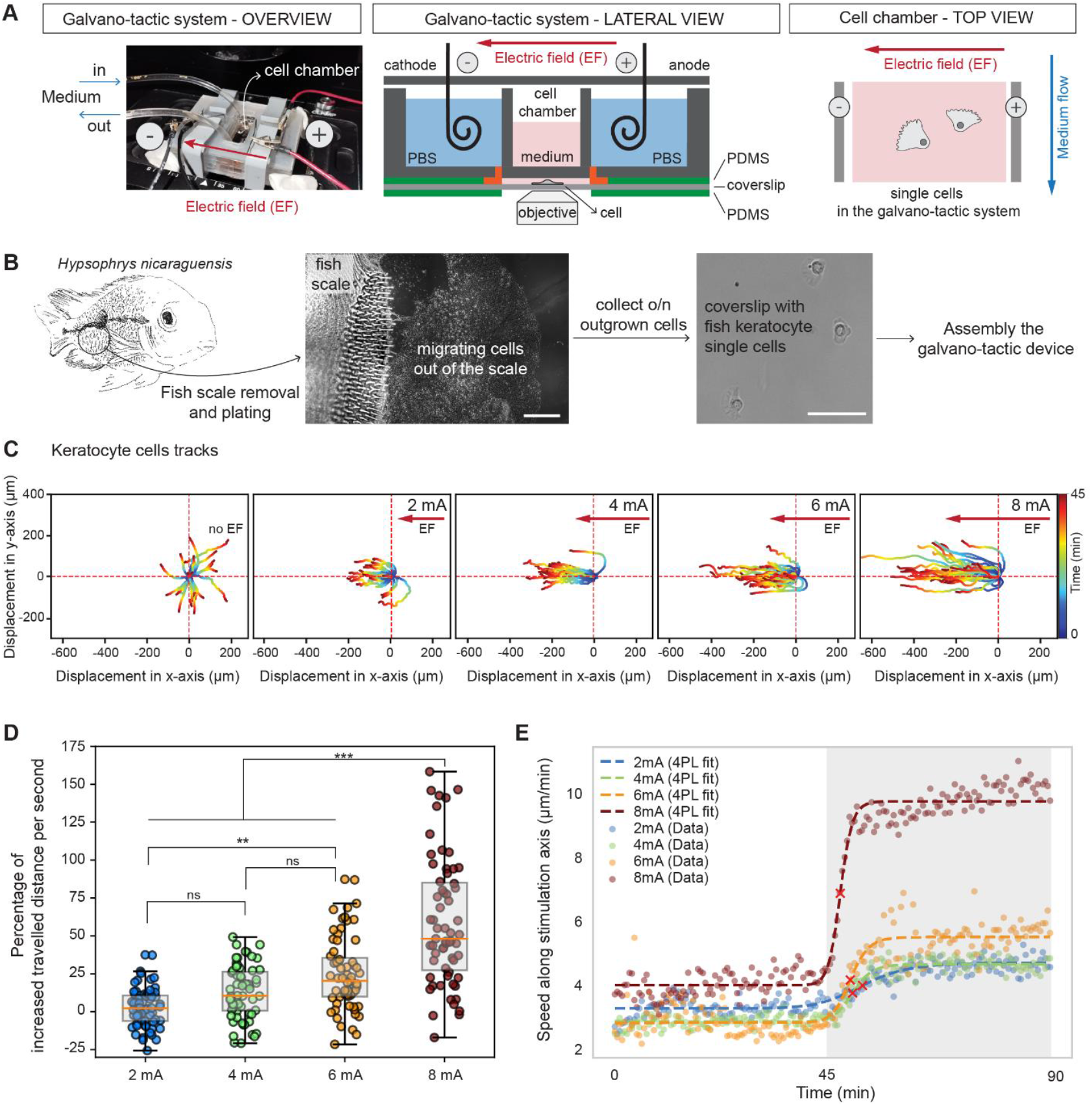
Electric field intensity regulates keratocyte migration direction and speed. **(A)** Illustration of the SCHEEPDOG^16^ (Cohen 2020) galvano-tactic system (overview, lateral and top view) for investigating electrotaxis in keratocyte cells. **(B)** Schematic representation of the experimental paradigm for isolation and electrical stimulation of fish keratocyte cells. **(C)** Displacement of keratocyte cell trajectories over a 45-minute imaging period normalized to a common origin, in control condition (no electrical field), and upon electrical stimulation at different electric field intensities (2mA, 4mA, 6mA, 8mA). Color-coding indicates time progression from dark blue (start) to dark red (end), as shown on the vertical scale bar. **(D)** Percentage of increased travelled distance per second for cells over the 45-minute period upon electrical stimulation at given intensities (2mA, 4mA, 6mA, 8mA) after 45 minutes of no stimulation. **(E)** Sigmoid fit of the average cell migration speed along the stimulation axis over time under different electrical stimulation intensities (Control, 2mA, 4mA, 6mA, and 8mA). Different stimulation intensities are color-coded. Data points are median values across cells at each time. The white section shows no stimulation period and shadowed section shows stimulation period. Red Cross represents the half time to reach maximum speed over stimulation time. Data information: Single tracks (C) and dots (D, E) correspond to single cell measures. Box plots represent the interquartile range (IQR), orange line correspond to median value, and whiskers extend to the extreme data points that are not considered outliers. Statistics: one-way ANOVA followed by Tukey’s multiple comparison post hoc test (D). Abbreviations: EF, electric field; ns, not significant. Scale bar: 500 μm (B, fish scale), 100 μm (B, single cells).

In our setup, increasing the stimulation current leads to a proportional increase in the applied electric field, assuming constant conductivity and fixed electrode geometry. Since the electric field strength (E) is determined by the applied voltage (V) over the distance between electrodes (d), and the current (I) is related to the voltage via Ohm’s law (V = IR), higher current intensities result in stronger electric fields under constant resistance conditions.

Interestingly, increasing the intensity of the electric field had a direct impact on the displacement of the cells along the direction of stimulation (Fig.1C). Regardless of stimulation intensity (2, 4, 6 and 8mA), keratocytes consistently exhibited directed migration aligned with the electric field (Supplementary movies 1-4). However, increasing the electric field intensity resulted in a marked enhancement of the travelled distance. Cells not only migrated more persistently along the field direction, but their overall displacement increased proportionally with field intensity. At the highest intensity tested (8 mA), the total travelled distance at the unit of time was much more than that observed under control conditions. The greater overall distance travelled by cells suggests an increase in their migration speed. To better show this increase in the amount of travelled distance, we have measured the velocity of each single cell before and during stimulation and then we have measured the percentage of increase in their velocity (Fig.1D). As we can see, at higher stimulation intensities cells show more increase in their velocity. To quantify the increase in speed along the stimulation axis, we measured the average speed of cells during the stimulation period and compared it to their average speed before stimulation (Fig.1E).

This approach enabled to study the dynamic changes in cell migratory behavior, assessing how cell speed evolved in response to increasing stimulation intensities. Under control conditions, cells maintain a relatively consistent speed over time (Fig.1E, first 45 minutes of all conditions). When a 2mA electrical stimulation is applied, cells gradually increase their speed, reaching 50% of the maximum speed after approximately 6 minutes. However, at the highest intensity tested (8mA), we observe an increase in speed at the onset of stimulation, followed by a slower increase. This rapid increase starts at the onset of stimulation. Notably, at 8mA, cells reach 50% of their maximum speed in around 90 seconds, following the initial sharp increase phase (Fig.1E). These observations highlight several key trends: first, at higher intensities, especially at 8mA, cells experience an immediate and sharp increase from the very beginning of the stimulation, followed by a gradual rise. Second, as the stimulation intensity increases, cells reach 50% of their maximum speed in progressively shorter timeframes (Details in supplementary Fig.1). Ultimately, regardless of the stimulation intensity, after a while, cells reach a maximum capacity in their velocity and maintain this velocity over time.

### Stronger electric fields accelerate cell alignment independently of migration distance

The pronounced increase in migration velocity across different stimulation intensities compelled us to further dissect the intricate dynamics of cellular response to electrical stimulation. Prior to stimulation, cells migrate in random directions, with subpopulations perpendicular to or even opposite to the field. However, upon stimulation, they progressively reorient and align with the electric field vector (Fig.2A). As higher faster migration is driven by stimulation intensities, this transition in speed raises a fundamental question: do cells also achieve alignment more rapidly in response to stimulation intensities? Alternatively, does alignment itself act as a prerequisite for enhanced motility? Exploring the relationship between speed and alignment can enhance our understanding of the mechanisms driving electrotactic behavior.

**Figure 2.**
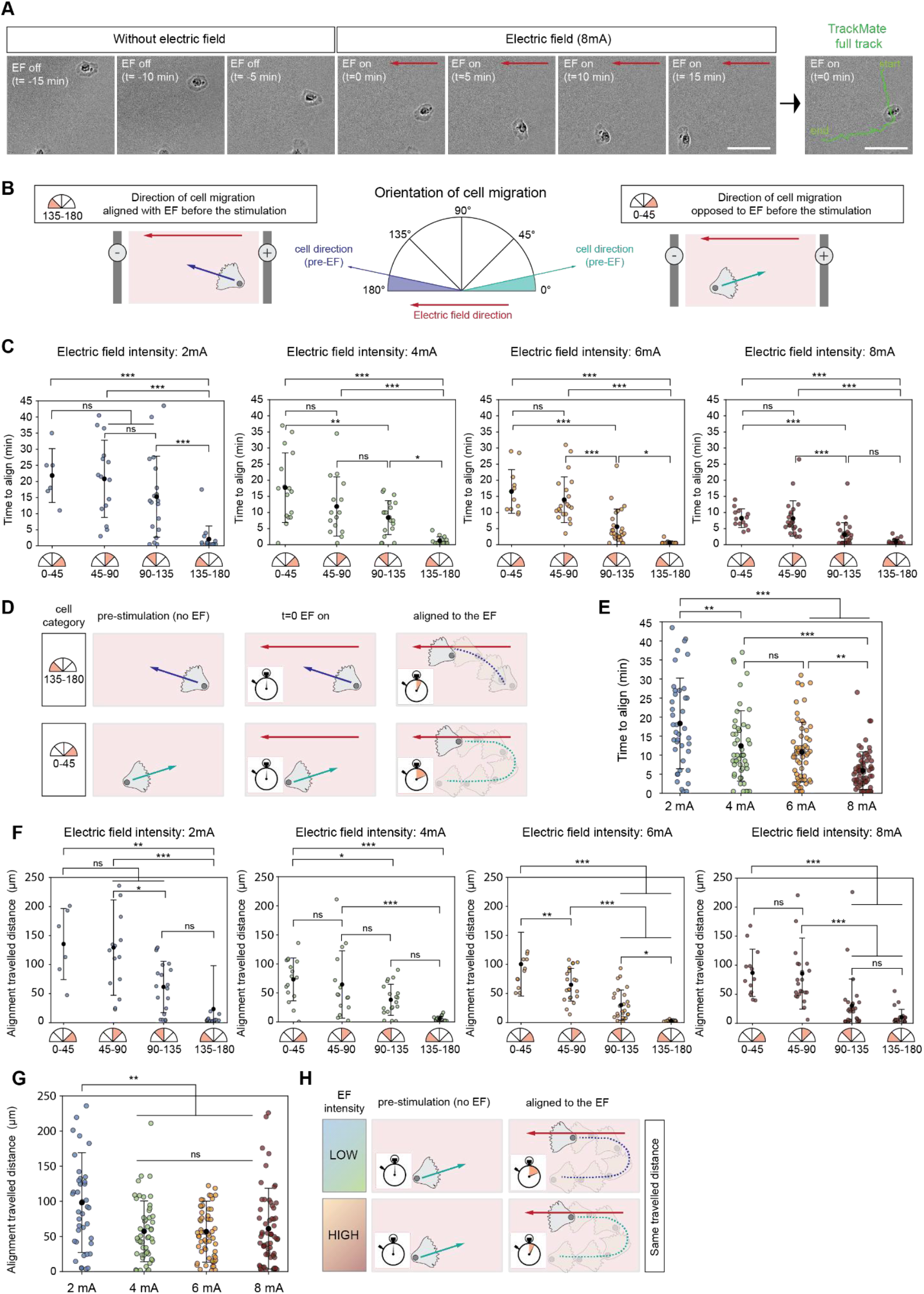
Stronger electric fields accelerate cell alignment independently of migration distance. **(A)** A single cell making a turn after applying the electric field at 8mA. Scale bar is 50 um **(B)** Schematic of analysis pipeline considering the initial orientation of cell migration (half-circle is divided into four 45° angular bins) in the process of alignment to the electric field. **(C)** Alignment time to electric field at different intensity based on initial orientation of cell migration (0°-45°, 45-90°, 90°-135°, 135°-180°). Colored sections in the half-circle diagrams represent the initial orientation bins of cell trajectories just before the onset of electrical stimulation. **(D)** Schematic representation of migration alignment to the electric field for cells initially moving in the EF direction (135°-180°) or opposite to it (0°-45°). **(E)** Mean alignment time across different stimulation intensities for collection of 0°-45°, 45-90° and 90°-135° groups. **(F)** Alignment travelled distance for cells grouped into four initial orientation bins based on their direction of migration prior to stimulation. **(G)** Mean alignment travelled distance across different stimulation intensities for collection of 0°-45°, 45-90° and 90°-135° groups. **(H)** Schematic representation of migration alignment to the electric field at different intensities. Data information: Swarm plot showing the data across different groups: Each dot represents an individual measurement. Mean ± standard deviation is overlaid as black circles with error bars. (C, E, F, G). Statistics: one-way ANOVA followed by Tukey’s multiple comparison post hoc test (C, E, F, G).

To investigate whether the initial migration direction influences the alignment process, we categorized trajectories into four angular bins, each spanning 45 degrees, based on their direction before stimulation (Fig.2B). Given that cells do not migrate along a perfectly straight path, we introduce the concept of an alignment cone, which represents a zone, that we defined arbitrarily, within which a trajectory is considered aligned to the electric field if it remains with minimal deviation of + or minus 36°. This classification enables the evaluation of how the initial migration direction affects subsequent alignment behavior in response to stimulation (Fig.2B). The first key parameter we measured is the alignment time, seeking to determine whether a cell’s initial migration direction influences the time required to align with the electric field. Across all stimulation intensities, a clear trend emerged: the smaller the initial deviation from the final alignment direction, the faster the alignment occurred (Fig.2C). Cells that initially migrated in the opposite direction of the field (0–45 degrees) required the longest time to reorient (Fig.2C, D), suggesting that alignment with the electric field does not occur through a simple reversal of polarity. In addition to the cell’s initial orientation, the intensity of the electric field plays a key role in regulating the speed of the alignment process. Indeed, the overall alignment time for cells that are not already aligned and need to make a turn to align with the direction of stimulation (0–45°, 45–90°, and 90–135° groups) decreases with increasing electric field intensity (Fig.2E), which can be attributed to the increased speed.

We next examined the distance traveled by cells from the onset of stimulation to the moment they achieved alignment, providing deeper insight into how migration dynamics adapt to varying stimulation intensities. A consistent pattern emerged across all conditions: the distance required for alignment mirrors the trend observed in alignment time. Specifically, cells initially migrating in the opposite direction of the electric field traveled a greater distance before aligning, whereas those with a smaller initial angle deviation required a shorter distance (Fig.2F) to achieve alignment. However, for cells that are not already aligned (0–45°, 45–90°, and 90–135°groups) and need to make a turn to align with the direction of stimulation, the average total travelled distance to reach alignment remained relatively consistent across all stimulation intensities (Fig.2G).

Overall, our data suggest that the stronger electric fields reduce the time required for cells to align their movement with the field, without significantly affecting the total distance traveled (Fig.2H). The electric field thus affects only the speed, not the trajectory.

### Trajectory shape remains unchanged while angular velocity drives faster alignment

To further explore how cells respond to varying stimulation intensities, we examined the shape of their trajectories during the alignment process (Fig.3A). Rather than focusing solely on the time or distance required for alignment, we sought to determine whether cells gradually reorient through smooth, continuous turns or whether they undergo abrupt, corrective changes in their migration paths.

**Figure 3.**
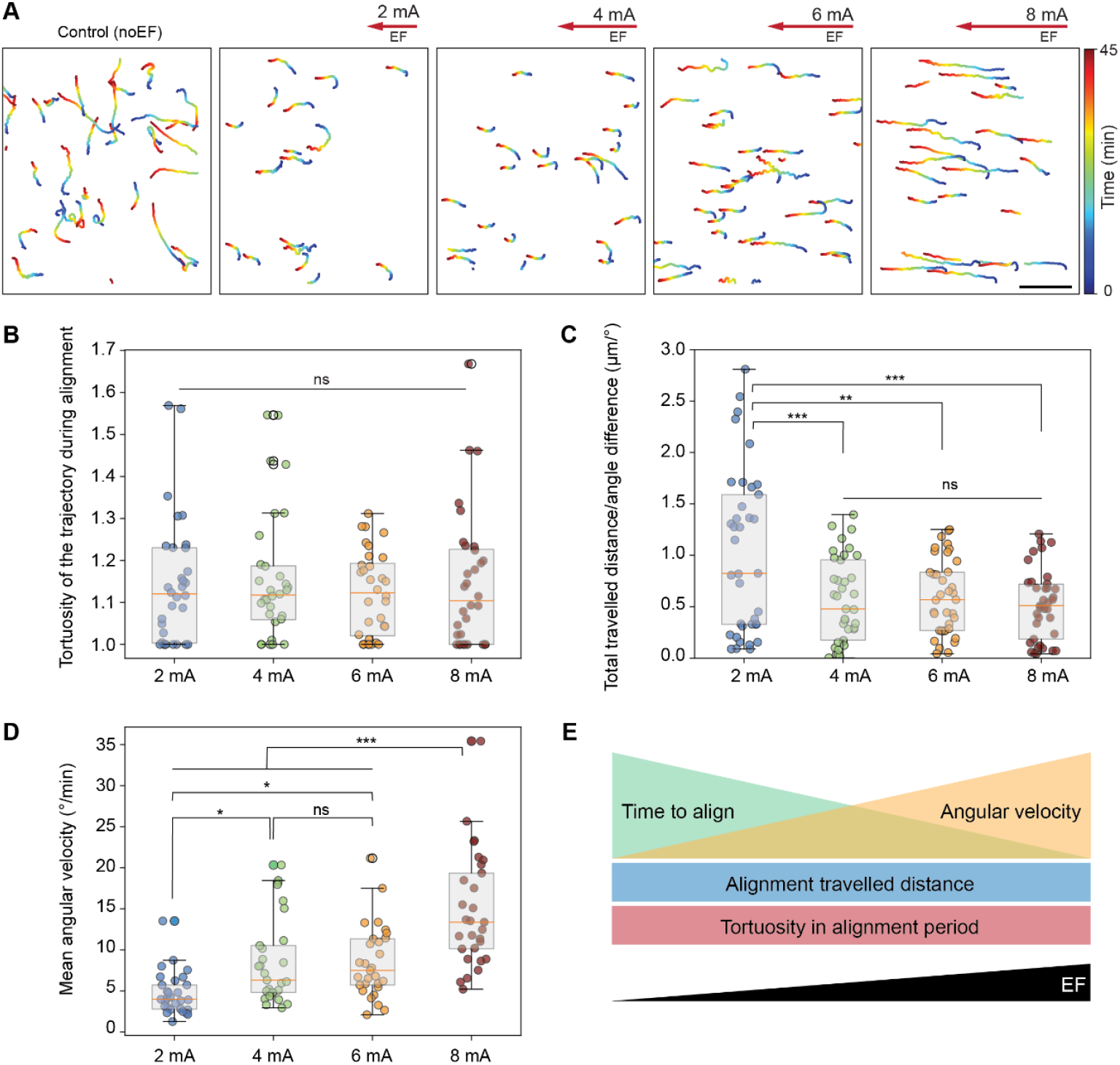
Trajectory shape remains unchanged while angular velocity drives faster alignment. **(A)** Trajectories of control cells and turning cells over 45 minutes of stimulation at different stimulation intensities (2mA, 4mA, 6mA, 8mA). **(B)** Average tortuosity values for each trajectory from the onset of stimulation to the point of alignment with the electric field. **(C)** Angular Path Ratio for each trajectory from the onset of stimulation to the point of alignment, incorporating the total travelled distance, initial angle prior to stimulation, and final angle at alignment. **(D)** Mean angular velocity from the onset of stimulation to the point of alignment with the electric field at each stimulation intensity. **(E)** Schematic represents cell alignment behavior under increasing stimulation intensities. Data information: Box plots represent the interquartile range (IQR), orange line corresponds to median value and whiskers extend to the extreme data points that are not considered outliers (B-D). Statistics: one-way ANOVA followed by Tukey’s multiple comparison post hoc test (B-D). Each circle represents one measurement and is color-coded according to the stimulation intensity. Scale bar for the trajectories (A): 500 μm.

To this end, we analyze the tortuosity of cell trajectories, which provides insight into the shape of the migration path. Tortuosity is defined as the ratio of the total traveled distance to the net displacement, with higher values indicating more winding trajectories and lower values reflecting more direct movement.

In this analysis, we measured the tortuosity of each trajectory from the onset of stimulation to the point at which the cells reached alignment. The average tortuosity values across different stimulation intensities did not exhibit significant differences (Fig.3B). However, cells that initially migrated in the direction opposite to the electric field displayed slightly higher tortuosity across all stimulation conditions.

To further characterize trajectory shape, we assessed curvature by calculating the Angular Path Ratio, which integrates the total distance traveled, the initial orientation before stimulation, and the final orientation at alignment. This metric captures the overall bending of the migration path during the alignment phase (Fig.3C).

Interestingly, the Angular Path Ratio of the trajectories showed no consistent trend with increasing electric field intensity (Fig.3C). In other words, despite the effect of electric field strength on alignment time, trajectory curvature remained relatively constant. This indicates that, while the alignment time was influenced by the electric field intensity, the shape of the alignment paths remained relatively unaffected by varying field strengths.

If the total travelled distance by the cells to achieve alignment remains unaffected by the intensity of the electric field, and if the quality of tortuosity and curvature over the alignment period shows no significant changes across different conditions, how are cells aligning faster at higher intensities? Since the overall migration paths and shapes of the cells remain similar, another potential factor influencing faster alignment is the rate at which they rotate toward the direction of the electric field. To investigate this, we measured the mean angular velocity of the cells during the alignment period. We observed that with increasing stimulation intensity, the angular velocity of the cells during the alignment period also increases (Fig.3D).

Altogether, our findings indicate that while increasing the intensity of the electric field reduces the alignment time and increases the angular velocity of the cells during alignment, it does not significantly affect the total distance travelled by the cells or alter the shape of their trajectories throughout the alignment process (Fig.3E).

## Discussion

Understanding cellular migration is essential for studying many biological processes. One well-known example is wound healing, where epithelial cells migrate to the margins of a wound to facilitate tissue repair^39^.

Fish keratocyte cells participate in skin wound healing process^40^. These cells are equipped with a lunate lamellipodium at the front and a spindle-shaped structure at the rear^41^. Actin polymerization drives the forward movement at the leading edge^42,43^, while myosin II contraction at the rear helps retract the back of the cell^42^ and enables keratocytes to turn^44^ and migrate efficiently.

In this study, we explored how varying electrical stimulation intensities affect the migration speed, alignment, and trajectory shape of keratocytes, which are model cells for studying electrotaxis^45^. Our findings reveal several important insights into the cellular dynamics under different field intensities.

In response to electrical stimulation, many different cells generally migrate aligned with the direction of the electric field toward the cathode. This migration has been studied at the tissue-level^15,16,46^or in single-cells^21^ like the work presented here.

In line with previous work, we observed that migrating keratocytes align with the direction of the electric field over the course of the stimulation. We showed that this behavior is consistent across all the tested electric field intensities, including at low intensity (i.e. 2 mA).

Cells stimulated at 8mA travelled approximately twice the distance compared to control conditions, highlighting the clear influence of increased field intensity on cellular migration.

The speed of the cells increased over the course of the stimulation (Supplementary Fig.2A), and when comparing the rate of this speed increase in individual cell trajectories to the no stimulation condition for the same cells, we found that the rate of increase was directly influenced by the intensity of the electric field (Fig.1E). Specifically, cells exposed to higher stimulation intensities exhibited a more pronounced and rapid increase in speed compared to those under lower intensities or control conditions. This highlights the direct relationship between the strength of the electric field and the acceleration of cell migration.

Interestingly, at higher stimulation intensities, the increase in cell speed and directionality persisted to some extent even after the stimulation was ceased (Supplementary Fig 2B, C) which is in accordance with previous studies^21^. This suggests that stronger electric fields may induce a lasting effect on the cell’s migratory behavior, potentially altering their internal dynamics or motility mechanisms even after the external stimulus is removed. This finding raises intriguing questions about the long-term impact of electrical fields on cellular migration and warrants further investigation into the mechanisms underlying this persistent effect. This point has already been discussed by Theriot’s lab where they claim that the time for a cell to lose directional migration after field removal depends on the viscosity of the fluid and the previous degree of cell polarization.

A detailed analysis of speed shows that initially, cells experience a rapid increase in speed at the onset of stimulation and then they reach an almost constant maximum speed. Notably, this maximum speed is consistently higher than the speed observed without stimulation. Measuring the speed over stimulation time reveals that cells respond almost immediately to electrical stimulation. Additionally, the time required to reach the maximum speed decreases as the stimulation intensity increases. This suggests that higher electric field intensities facilitate a faster cellular response, allowing cells to reach their maximum migration speed more quickly.

Importantly, as we have also discussed previously, the overall increase in cell speed is likely influenced, by heating resulting from higher electric field intensities, which can elevate the local temperature and thereby enhance cellular motility. However, as shown in Fig.1E, this increase in speed is predominantly oriented along the stimulation axis, indicating that the directionality and alignment of movement are governed by the electric field itself, rather than by thermal effects.

To further understand the effect of stimulation magnitude on cell behavior, we investigated the alignment process during stimulation. Cells with a smaller initial angle relative to the electric field vector exhibited shorter alignment times (Fig.2C). This trend was consistent across all applied stimulation intensities. Furthermore, we observed that increasing the stimulation intensity resulted in a further reduction in average alignment time regardless of the initial direction (Fig.2E). Overall, our findings indicate that stronger electrical fields accelerate the alignment of cells with the electric field direction.

Comparing the speed of cells over stimulation time with their average alignment time reveals a key observation: across all conditions, the time required for cells to reach 50 percent of their maximum speed is consistently shorter than the time needed for full alignment. This suggests that cells begin accelerating in response to the electric field before they completely align with its direction. The shorter time to reach maximum speed indicates that the increase in migration velocity is an immediate response to stimulation, whereas alignment occurs more gradually as the cells adjust their direction over time.

Similar to alignment time, cells with smaller angular deviation, align with smaller travelled distance at all stimulation intensities (Fig.2F), but regardless of the initial direction, the average distance travelled by cells before achieving alignment remains consistent across all conditions (Fig.2G).

However, analyzing the trajectories of cells during alignment, along with the greater travelled distances required for cells initially moving opposite to the electric field, reveals that cells usually make a turn to align to the direction of the field and as reported before^33,44^, only in very rare occasions they flip and repolarize. This leads us to question how cells adjust their migration paths during alignment. Do they follow a smooth, gradual curve or do they take a more abrupt, corrective turn? To investigate this, we next examined the shape of cell trajectories throughout the alignment process.

Measuring the tortuosity of the alignment path and the curvature of trajectories over the alignment period shows that increasing the stimulation intensity does not significantly affect the overall shape of the trajectories. However, the angular velocity of the cells increases significantly with higher stimulation intensities, suggesting that while the trajectory shape remains unchanged, the rate at which cells reorient is enhanced (Fig3E).

In summary, our findings demonstrate that keratocyte cells exhibit a robust response to electrical stimulation, regardless of the intensity of the applied electric field. The observation that increased motility precedes directional alignment suggests that the primary effect of the electric field acts on the cell’s motility machinery rather than directly on its guidance mechanisms. Importantly, cells that initially migrated against the direction of the electric field did not simply undergo repolarization. Instead, the prolonged time and greater distance required for these cells to reorient reinforce the idea that re-alignment involves more complex processes than a mere polarity switch.

Moreover, our data indicate that the mode of cellular turning is not modulated by the strength of the electric field. This suggests that the turning behavior is governed intrinsic properties of the cytoskeleton, which may operate near a functional threshold that limits its ability to respond to external cues such as electric stimulation. The forward translocation of the cell body and retrograde flow within the lamellipodium are driven by coordinated interplay of cytoskeletal components, including actin– myosin network within the lamellipodium^42^, actin polymerization at the leading edge^47,48^ and myosin II– mediated contraction at the rear^49^.

What are the mechanisms underlying the turning behavior observed during electrotaxis? Does the electric field reorganize the actin cytoskeleton, influence adhesion dynamics, or activate associated signaling pathways? It is also possible that a combination of these factors contributes to the observed cellular response. One plausible scenario is that actin-binding proteins, which regulate cytoskeletal organization, are indirectly affected by the electric field, leading to changes in migration dynamics. These questions remain open and represent important directions for future research into the biophysical and molecular mechanisms of electrically guided cell migration.

## Material and methods

### Experimental model and cell culture

The primary culture medium consists of Leibovitz’s L-15 Medium (Gibco™, Thermo Fisher Scientific, Cat. No. 11415064) supplemented with 10% fetal bovine serum (FBS) and penicillin-streptomycin. Then mix 10mL of primary culture medium, 300µL of Hepes buffer pH=7.4, 3mL of deionized water to prepare what we refer to as keratocyte medium.

Keratocyte cells were harvested from the scales of the Central American cichlid fish *Hypsophrys Nicaraguensis*. Without sacrificing the fish, scales were gently removed from the flank of a fish and rinsed with keratocyte media at room temperature. Scales were then placed in small 35mm sterile petri dishes and sandwiched with 22mm glass coverslips. Incubated overnight at room temperature to promote adherence. Keratocytes were cultured in the dark at room temperature for 12 h. Individual keratocytes were then dissociated by incubating the epithelial tissue with trypsin and EGTA for 5 min and resuspended in L-15 Leibovitz complete medium. Cell suspension was carefully pipetted onto the glass coverslips and allowed to settle for 70 minutes. After this incubation period, sufficient culture medium was added to fill the dish, preparing it for assembly onto the experimental device.

To avoid physical contact between cells and ensuring the study of single cell galvanotactic behaviour, we maintained a nearly homogeneous cell density throughout all experiments, at approximately 20 cells/mm^2^ (Supplementary Figure 3).

### Hoechst staining

Nuclei were stained using Hoechst dye (Hoechst 33342, Thermo Fisher Scientific) to allow visualization under the DAPI fluorescence channel. Cells were incubated with the dye for 10 minutes at room temperature, followed by gentle washing with PBS prior to imaging.

For electrical stimulation, the SCHEEPDOG system, provided by CohenLab, was used^16^. A Keithley source meter (Keithley 2400/2450, Tektronix) supplied current to the stimulation electrode pairs, while a USB oscilloscope (Analog Discovery 2, Digilent Inc.) measured the voltage across the pair of recording probes.

Custom MATLAB script from Cohen Lab, was used to control the instruments, drive a set current, and measure the resultant voltage across the stimulation chambers. Proportional feedback control was implemented to adjust the output current, ensuring the target field strength and direction were maintained.

In our experiments, we applied currents in the range of 2–8 mA. Using voltage probes installed in the chamber, we recorded corresponding electric field strengths ranging from 0.2 to 7 V/cm, corresponding to applied currents between 2 and 8 mA. Given the geometry of the chamber, the current density J can be expressed as:

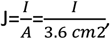

where *I* is the applied current and *A* is the cross-sectional area of the chamber (3.6 cm^2^).

### Pumping system

A peristaltic pump (Instech Laboratories) was used to continuously circulate fresh media through the cell chamber at a flow rate of 2.5 ml per hour.

### Microscopy

Cells were observed on an inverted Nikon Ti-E2 microscope with an Orca Flash 4.0 sCMOS camera (Hamamatsu). A Nikon ×20 objective was used and for the experiments NIS-Elements software (Nikon Instruments Inc., Tokyo, Japan) was used for controlling the microscope and camera during image acquisition.

### Trajectory analysis

For trajectory analysis, we utilized the Fiji distribution of ImageJ, specifically the TrackMate plugin^50^, to track and analyze cell movement. This tool enabled automated tracking of individual cells over time during live imaging. From the extracted trajectory data, we calculated key migration parameters including instantaneous and average speed, total travelled distance, and tortuosity. It is important to note that for all experiments, we specifically limited our imaging and analysis to the middle portion of the chamber, where the field distribution is more likely to be well characterized and uniform (which has been quantified and shown by Cohen’s lab^16^.

### Calculations

*Speed calculated between two consecutive time points as following:

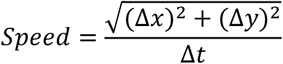

Δ*x and* Δ*y* are the difference in the X and Y position of the cell between two consecutive time points and Δ*t* shows the time difference between the two points.

To see how the speed changes along the stimulation axis, we have plotted the average speed of several cells over time then we have used a 4PL sigmoid curve to model how speed changes over time after stimulation. The curve helps describe how quickly and how strongly each group responds to stimulation. It provides interpretable measures like how long it takes the speed to start increasing and how steeply it increases.

To do this, we first applied a smoothing function (spline smoothing) to reduce noise in the speed data for each condition. Then we fit data to 4-parameter logistic (4PL) curve:

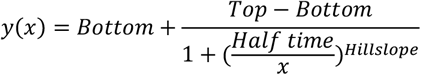

where, Bottom is minimum value of the response, and Top is maximum value of the response. Half-time is the value of at which the function reaches halfway between Bottom and Top. Finally, Hillslope describes the steepness of the transition between Bottom and Top.

*The alignment point is determined as the first moment during the stimulation period when the direction of movement falls within a predefined angular range—in this case, between 144° and 180° (This cone has been defined arbitrarily). This indicates the point at which the movement begins to align with the target region.

Since trajectories are not always perfectly linear, a deviation threshold is applied: if the proportion of direction segments falling outside the target angle range is less than or equal to 30%, the motion is considered aligned.

In summary, the alignment point is defined as the earliest time during stimulation when the movement direction enters the specified angular range and remains mostly within it.

To calculate the alignment travel distance, the total distance moved from the start of stimulation to the alignment point is measured. To determine the initial trajectory angle, we used the positions at four consecutive time points immediately before the onset of stimulation. The angle was calculated between the first and last of these four positions, providing a measure of the overall direction of movement during this brief pre-stimulation window. This angle was computed relative to the x-axis, using the arctangent of the displacement vector. Trajectories were then grouped into four angular ranges: 0°–45°, 45°–90°, 90°– 135°, 135°–180°. This method captures the predominant direction of movement just before stimulation begins. To assess how much each trajectory deviated from a straight path during the stimulation period, we calculated tortuosity at the point of alignment. Tortuosity was defined as the ratio of the total travelled distance from the start of stimulation to the alignment time, divided by the straight-line distance between those two points. A value close to 1 indicates a nearly straight trajectory, while higher values represent more winding or irregular paths. Additionally, based on the initial movement angle before stimulation, each trajectory was categorized into one of four angular groups (0°–45°, 45°–90°, 90°–135°, and 135°– 180°) to analyze directional differences in path characteristics.

### Angular Path Ratio calculation

To assess how efficiently movement occurred relative to changes in direction, we calculated an Angular Path Ratio, defined as the ratio of total travelled distance from start of stimulation to the alignment time to the angular change in trajectory between the beginning of stimulation and alignment time. This metric reflects how much distance was covered per unit change in movement direction.

To quantify how rapidly trajectories reoriented after stimulation began, we calculated the mean angular velocity between stimulation onset and the time of first alignment with the alignment cone. Angular velocity was computed as the rate of change in direction (in degrees per unit time), based on movement angles unwrapped to account for circular discontinuities. The average of these values represents the mean angular velocity for each trajectory.

### Statistics

Each box plot represents the interquartile range (IQR), orange line corresponds to median value, and whiskers extend to the extreme data points that are not considered outliers. Swarm plots showing the data across each condition: Each dot represents an individual measurement. Mean ± standard deviation is overlaid as black circles with error bars. For the statistical analysis, we performed one-way ANOVA followed by Tukey’s multiple comparison post hoc test. For all tests, the significance level was α = 0.05 (\**P* < 0.05, \*\**P* < 0.01, \*\*\**P* < 0.001).

## Author contributions

M.B., N.P., E.C. D.C., G.S, M.L. and T.B. conceived the study and designed the experiments. N.P performed experiments. N.P,E.C,G.C. analyzed the data. N.P,E.C and M.B. wrote the manuscript with feedback from all authors. M.B., S.G. and D.C. supervised the project.

## Data availability

Because of the large file size, the datasets generated and/or analyzed during the current study are available from the corresponding author on request. A response will be provided in less than 2 weeks.

## Acknowledgements

M.B. acknowledges financial support from the ANR PlatForMech project, grant ANR-18-CE14–0037-02 and the ANR Inter-s-cal project, grant ANR-21-CE13–0042-02 of the French Agence Nationale de la Recherche (ANR). G.C. acknowledges financial support from the ANR SupraWaves project, grant ANR-19-CE13-0028 of the French ANR. T.B. and S.d.B. acknowledge funding from CNRS grants (PEPS CNRS-INSIS 2021, Lumière Visible et Vie 2022).

