## Supplementary Figures for "Electric field intensity modulates keratocyte migration without altering turning dynamics"

(1) Univ. Grenoble Alpes, CNRS, LIPhy, 38000 Grenoble, France

(2) Interdisciplinary institute for neuroscience (IINS), University of Bordeaux, UMR 5297, F-33000 Bordeaux, France

(3) Department of Mechanical and Aerospace Engineering, Princeton University, Princeton, NJ 08544, USA.

(4) SYMBIOSE Lab, CIRMAP, Research Institute for Biosciences, University of Mons, Mons, Belgium

| Condition | Half time | HillSlope |
|---|---|---|
| 2mA | 6.130 | 10.901 |
| 4mA | 4.115 | 25.176 |
| 6mA | 3.632 | 26.133 |
| 8mA | 1.590 | 38.057 |

**Supplementary Figure 1.** The table shows the values for half time to reach the maximum speed along the field axis and the steepness of the transition between Bottom and Top (Hillslope) at different stimulation intensities (Condition).

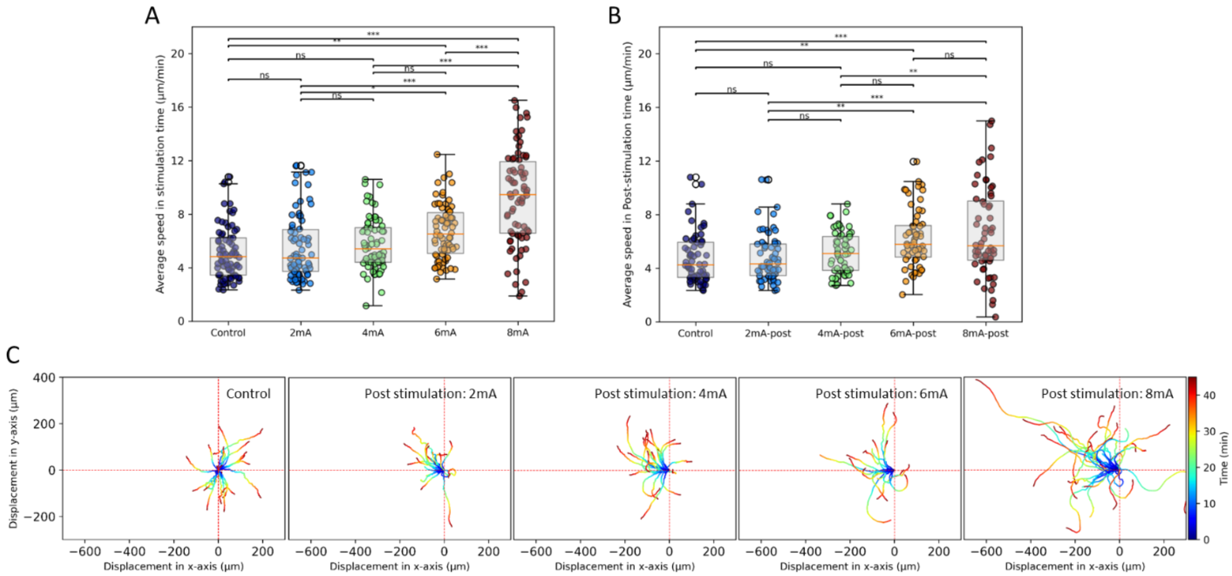


**Supplementary Figure 2. (A)** Average speed of cells over the 45-minute period electrical stimulation at given intensities (2mA, 4mA, 6mA, 8mA). **(B)** Average speed of cells after the 45-minute period electrical stimulation at given intensities (2mA, 4mA, 6mA, 8mA). **(C)** Displacement of keratocyte cell trajectories (for 45 minutes) after a stimulation period normalized to a common origin, in control condition (no electrical field), and after ceasing electrical stimulation at different electric field intensities (2mA, 4mA, 6mA, 8mA).

Data information: Single tracks and dots (A, B) correspond to single cell measures (For A: EF=2mA, 4mA, 6mA and 8mA and for B: Post-stimulation at 2, 4, 6 and 8mA). Box plots represents the interquartile range (IQR), orange line correspond to median value, and whiskers extend to the extreme data points that are not considered outliers. Statistics: one-way ANOVA followed by Tukey's multiple comparison post hoc test (A, B). Abbreviations: ns, not significant. Color-coding (C) indicates time progression from dark blue (start) to dark red (end), as shown on the vertical scale bar.

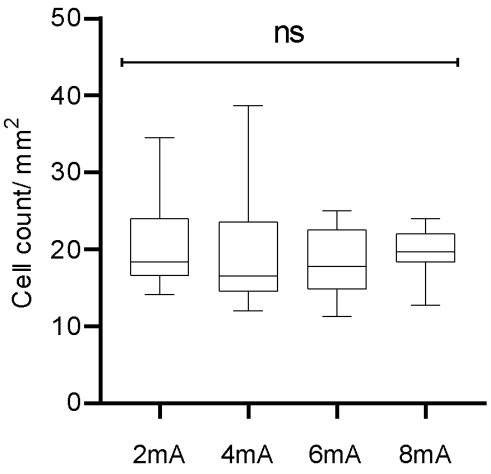


**Supplementary Figure 3.** Average number of the cells at each condition. Cell counts were obtained for each frame across multiple experiments conducted separately for each condition.
